# The interplay of demography and selection during maize domestication and expansion

**DOI:** 10.1101/114579

**Authors:** Li Wang, Timothy M. Beissinger, Anne Lorant, Claudia Ross-Ibarra, Jeffrey Ross-Ibarra, Matthew B. Hufford

## Abstract

The history of maize has been characterized by major demographic events including changes in population size associated with domestication and range expansion as well as gene flow with wild relatives. The interplay between demographic history and selection has shaped diversity across maize populations and genomes. Here, we investigate these processes based on high-depth resequencing data from 31 maize landraces spanning the pre-Columbian distribution of maize as well as four wild progenitor individuals (*Zea mays* ssp. *parviglumis*) from the Balsas River Valley in Mexico. Genome-wide demographic analyses reveal that maize domestication and spread resulted in pronounced declines in effective population size due to both a protracted bottleneck and serial founder effects, while, concurrently, *parviglumis* experienced population growth. The cost of maize domestication and spread was an increase in deleterious alleles in the domesticate relative to its wild progenitor. This cost is particularly pronounced in Andean maize, which appears to have experienced a more dramatic founder event when compared to other maize populations. Introgression from the wild teosinte *Zea mays* ssp. *mexicana* into maize in the highlands of Mexico and Guatemala is found found to decrease the prevalence of deleterious alleles, likely due to the higher long-term effective population size of wild maize. These findings underscore the strong interaction between historical demography and the efficiency of selection species- and genome-wide and suggest domesticated species with well-characterized histories may be particularly useful for understanding this interplay.

## 1 Introduction

Genomes are shaped over the course of their evolutionary history through a complex interaction of demography and selection. Neutral processes that comprise a species’ demographic history, such as stochastic changes in population size and migration events, influence both the pool of diversity upon which selection can act and its efficiency. Selection and genetic drift then jointly determine the fate of this diversity.

After the development of agriculture, both crops and humans have experienced profound demographic shifts that left clear signatures in genome-wide patterns of diversity [1, 2]. Early agriculturalists sampled a subset of the diversity present in crop wild relatives, resulting in an initial demographic bottleneck for many domesticates [3]. Subsequent to domestication, humans and their crops experienced a process of global expansion facilitated by the rise of agriculture [4]. In many cases expansion was accompanied by gene flow with close relatives, a demographic process that further altered patterns of diversity [5, 6].

Recent interest in the effects of demography on functional variation has led to a growing body of theory that is increasingly supported by empirical examples. To date, the relationship between demography and selection has been most thoroughly explored in the context of deleterious alleles. While theory suggests mutation load may be insensitive to demography over long periods [7, 8], empirical results are consistent with load being shaped by demography over shorter timescales [9, 10, 11, 12, 13]. For example, evidence in both plant and animal species has revealed increased mutation load in populations that have undergone recent, sudden declines in effective population size (*N*_*e*_) [10, 11, 12, 14] Similarly, in geographically expanding populations, repeated sub-sampling of diversity (*i.e.*, serial founder effects) can occur during migration away from a center of origin [15, 16], a phenomenon shown to have decreased genetic diversity and increased counts of deleterious alleles in human populations more distant from Africa [17, 18]. Finally, gene flow may also affect genome-wide patterns of deleterious variants, particularly when occurring between populations with starkly contrasting *N*_*e*_. For instance, during the Out-of-Africa migration, modern humans inter-mated with Neanderthals, a close relative with substantially lower *N*_*e*_ and higher mutation load [9]. The higher mutation load in Neanderthals presented a cost of gene flow, and subsequent purifying selection appears to have limited the amount of Neanderthal introgression near genes in the modern human genome [9, 19].

The domesticated plant maize *(Zea mays ssp. mays)* has a history of profound demographic shifts accompanied by selection for agronomic performance and adaptation to novel environments, making it an ideal system in which to study the interaction between demography and selection. Maize was domesticated in a narrow region of southwest Mexico from the wild plant teosinte *(Zea mays* ssp. *parviglumis*; [20, 21, 22]) and experienced an associated genetic bottleneck that removed a substantial proportion of the diversity found in its progenitor [23, 24]. Archaeological evidence suggests that after initial domestication, maize spread across the Americas, reaching the southwestern US by approximately 4,500 BP [25] and coastal South America as early as 6,700 BP [26]. Gene flow into maize from multiple teosinte species has been documented in geographical regions outside of its center of origin [5, 27]. To date, genetic studies of demography and selection in maize have primarily focused on initial domestication [28], only broadly considering the effects of subsequent population size change on diversity [2] and largely disregarding the spatial effects of geographic expansion and gene flow (but see [29]). Furthermore, the effect of maize demography on the prevalence of deleterious alleles has yet to receive in-depth attention.

Here, we investigate the genome-wide effects of demographic change in maize during domestication and subsequent expansion using high-depth resequencing data from a panel of maize landraces. We present evidence for a protracted domestication bottleneck, further loss of diversity during crop expansion, and gene flow between maize and its wild relatives outside of its center of origin. We then explore how this demographic history has shaped genome-wide patterns of deleterious alleles.

## 2 Results

### Maize population size change during domestication and expansion

We resequenced 31 open-pollinated maize landraces representing six geographical regions that span the pre-Colombian range of maize cultivation (Figure 1) as well as four wild *parviglumis* individuals from a single population located in the Balsas River Valley in Mexico. Median sequencing depth was 29X, with a range of 24-53X. Landrace accessions were selected to broadly reflect the diversity of maize in the Americas and to be representative of defined ecogeographic regions based on consultation with experts on landrace germplasm (Major Goodman, personal communication) and on descriptions in the “Races of Maize” handbooks [30].

**Figure 1:**
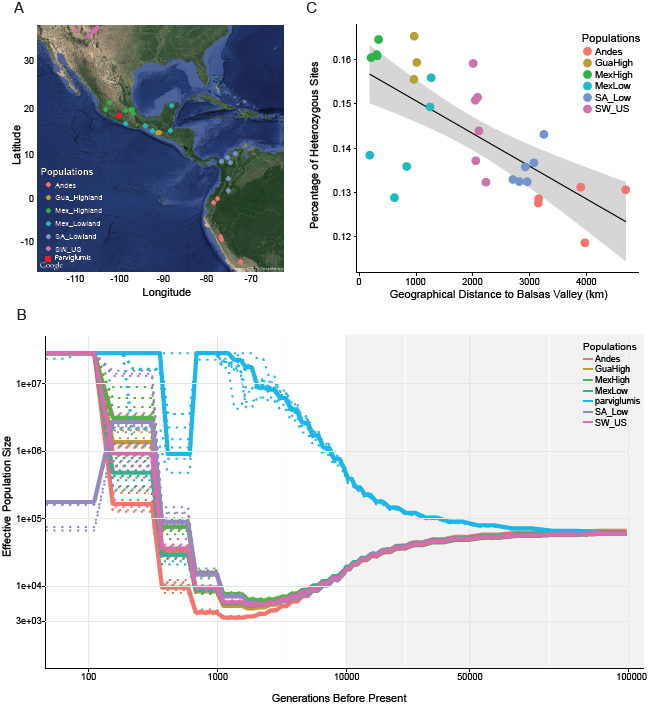
Maize domestication and expansion. A. Sampling locations. B. Estimates of effective population size over time. Dashed lines represent bootstrapping results. The x axis is *log* 10 scaled when time is less than 10,000 generations BP and linear when greater than 10,000 generations BP as indicated by the grey background. C. The percentage of polymorphic sites versus distance from the maize domestication center. Abbreviations for populations: GuaHigh, Guatemalan Highlands; MexHigh, Mexican Highlands; MexLow, Mexican Lowlands; SA_Low, South American Lowlands; SW_US, Southwestern US Highlands.

We first estimated historical changes in effective population size (*N*_*e*_) of maize and *parviglumis* using the multiple sequentially Markovian coalescent (MSMC) [31]. Consistent with archaeological evidence [21], we find that the demographic histories of the various maize populations begin to diverge from one another approximately 10,000 years before present (BP; Figure 1B). Surprisingly, our single population of *parviglumis* diverges from maize much earlier, around 75, 000 generations BP. All maize populations show a gradual decline in diversity concomitant with divergence from *parviglumis*, but the slope becomes more pronounced around the time of domestication. This period of declining Ne continues until the recent past (≈ 1,100 — 2,400 generations BP) and is followed by extremely rapid population growth, suggesting recovery from domestication post-dated expansion of maize across the Americas. In contrast to our results in maize, *parviglumis* shows an increase in *N*_*e*_ which also lasts until the recent past (≈ 1, 200 — 1, 800 generations BP). To determine if linked selection associated with domestication could bias estimates of *N*_*e*_ in maize (see [32]) we masked previously identified domestication candidates [24] and observed nearly identical results (Figure S1A).

One explanation for the prolonged population size reduction in maize following the onset of domestication would be repeated colonization bottlenecks during spread across the Americas. Genome-wide levels of heterozygosity across our maize samples are consistent with this idea, showing a strong negative correlation (*R*^2^ = 0.3636,*p* = 0.0004; Figure 1C) with distance from the center of maize domestication in the Balsas River Basin. To confirm this trend, we performed a similar analysis with a much larger sample of published genotyping data (n = 3520; Figure S1) [S3] and observed similar results.

While the gradual decrease in genetic diversity seen with distance from the Balsas indicates serial founder effects, our analyses also point to a more extreme founder event in the Andean highlands of South America. Andean landraces show a deeper bottleneck in our MSMC analysis (Figure 1), have the lowest overall diversity (Figure S2), and show both a distinct reduction of low frequency alleles and a greater proportion of derived homozygous alleles compared to other populations (Figure S2). To shed light on the timing of this extreme founder event, we assessed evidence for recent inbreeding. Inbreeding coefficients in Andean samples were quite low and not statistically different from other populations (all *F <* 0.002 and *p >* 0.05 based on a Wilcoxon test). Likewise, no significant differences could be found across populations in the number of runs of homozygosity (ROH) longer than 1*cM* (*p >* 0.05 in all cases, Wilcoxon test), further suggesting a lack of recent inbreeding. However, when ROH were limited to those shorter than 0.5*cM*, a length of homozygosity that would be associated with a more ancient founder event, Andean samples demonstrated greater cumulative ROH genome-wide compared to all (*p <* 0.05, Wilcoxon test) but the South American lowland population (*p* = 0.214, Wilcoxon test). Together, these lines of evidence are consistent with an unusually strong founder event during colonization of the Andes.

### Introgression from wild maize in highland populations

Adaptive introgression from the wild teosinte taxon *Zea mays ssp*. *mexicana* (hereafter, *mexicana*) has previously been observed in maize in the highlands of Mexico [5]. Our broad sampling allowed us to investigate whether introgressed *mexicana* haplotypes have spread to highland maize populations outside of Mexico, potentially playing a role in adaptation in other regions. In order to test this hypothesis, we calculated Patterson’s D statistic [34] across all maize populations. All individuals from both the Mexican and Guatemalan highlands exhibited highly significant evidence for shared ancestry with *mexicana* (Figure S5A). Maize from the southwestern US also showed more limited evidence of introgression, consistent with findings from ancient DNA suggesting this region was originally colonized by admixed maize from the highlands of Mexico [35]. In contrast, the distribution of z-scores for South American populations overlapped zero, providing no evidence for spread of *mexicana* haplotypes to this region.

We localized introgression to chromosomal regions through genome-wide calculation of the 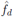 statistic [36]. Megabase-scale regions of introgression were identified in both Mexican and Guatemalan highland populations that correspond to those reported by [5] on chromosomes 4 and 6 (Figure 2; Figure S5). On chromosome 3, a large, previously unidentified region of introgression can be found in the Mexican and southwestern US highlands (Figure 2; Figure S5). This region overlaps a putative chromosomal inversion associated with flowering time in maize landraces [37] and in the maize nested association mapping population [38] and may be an example of *mexicana* contribution to modern maize lines.

**Figure 2:**
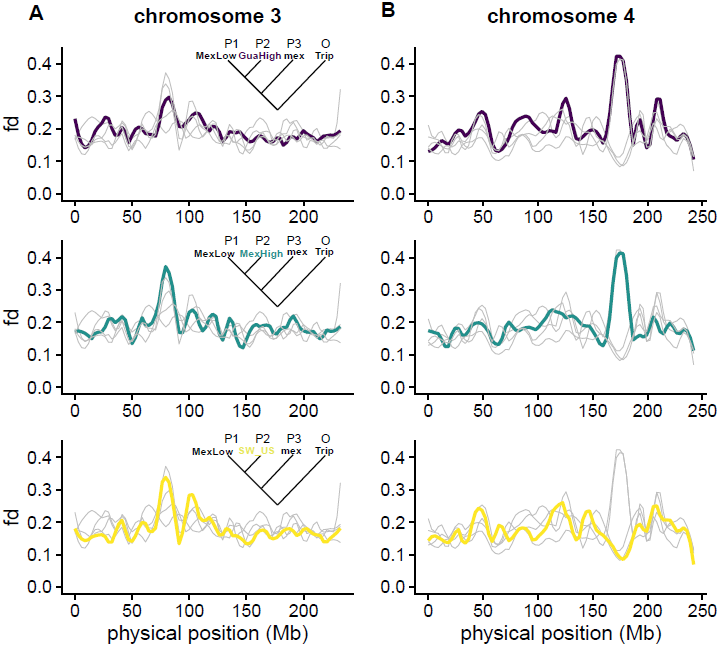
Introgression from *mexicana* into maize landraces. (A) chromosome 3 and (B) chromosome 4. The statistic 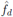 was calculated based on the tree, in which *P*_2_ is varied across highland and the South American lowland populations. Shown are results for the indicated three highland populations; other populations are drawn in grey. mex: *mexicana*; Trip: *Tripsacum*.

### The influence of demography on accumulation of deleterious alleles

Population-specific changes in historical *N*_*e*_ should influence the efficiency of purifying selection and alter genome-wide patterns of deleterious variants [10]. Introgression from a species with substantially different *N*_*e*_ may also influence the abundance and distribution of deleterious alleles in the genome [9]. Below we evaluate the effects of major demographic events during the pre-Colombian history of maize on patterns of deleterious alleles.

### Domestication and deleterious alleles

We first compared counts of deleterious alleles in Mexican lowland maize individuals to four *parviglumis* individuals from a single population in the Balsas River Valley. Maize from the Mexican lowlands has not experienced substantial introgression from wild relatives and is near the center of maize origin [22], and thus best reflects the effects of domestication alone. After identifying putatively deleterious mutations using Genomic Evolutionary Rate Profiling (GERP) [39], we calculated the number of derived deleterious alleles per genome under both an additive and a recessive model across four levels of mutation severity (see Methods for details). Maize showed significantly more deleterious alleles than teosinte under both additive (< 10% more; *p* = 0.0079, Wilcoxon test; Figure S6) and recessive (< 20 — 30% more; *p* = 0.0079; Figure 3) models across all categories (Figure S7). Additionally, maize contained 81% more fixed deleterious alleles than teosinte (48, 890 vs. 26, 947) and 3% fewer segregating deleterious alleles (464, 653 vs. 478, 594), effects expected under a domestication bottleneck (Figure 3; [7]). GERP load (GERP score x frequency of deleterious alleles), a more direct proxy of the individual-level burden of deleterious mutations, revealed a similar trend (additive model: maize median = 23.635, teosinte median = 22.791, *p* = 0.008, Wilcoxon test; recessive model: maize median = 14.922, teosinte median = 12.231, *p* = 0.008). Maize, like other domesticates [12, 14, 40, 41], thus appears to have a higher burden of deleterious alleles compared to its wild progenitor *parviglumis*.

**Figure 3:**
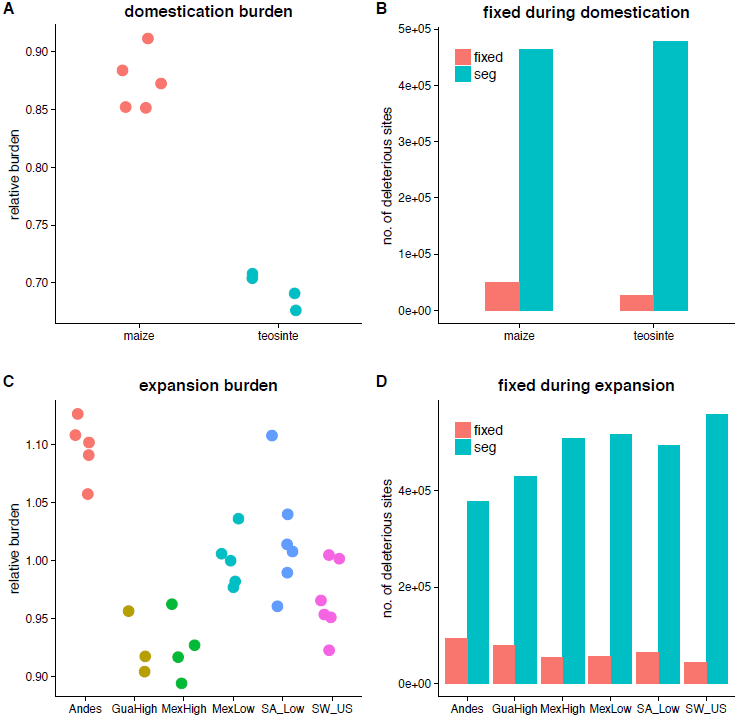
Burden of deleterious mutations during maize domestication and expansion. Comparison of counts of deleterious sites at the individual level (A) between *parviglumis* and maize and (C) among maize populations under a recessive model; comparison of fixed vs segregating (seg) deleterious sites at the population level (B) between *parviglumis* and maize and (D) among maize populations.

While the elevated burden we observe in maize relative to *parviglumis* may be driven primarily by the domestication bottleneck, positive selection on causal variants underlying domestication phenotypes may also fix nearby deleterious variants through genetic hitchhiking, which would result in a higher number of deleterious alleles in regions linked to domestication loci [40, 42]. To test this hypothesis, we first confirmed that 420 previously identified domestication candidates [24] showed evidence of selection in our data (Figure S8), and then assessed the distribution of deleterious alleles in and near (5*kb* upstream and downstream) these genes by calculating the number of deleterious alleles per base pair under both recessive and additive models. No significant difference was found in the prevalence of deleterious alleles near domestication and random sets of genes (Figure S9), suggesting the increased mutation burden of deleterious alleles in maize has been driven primarily by the genome-wide effects of the domestication bottleneck rather than linkage associated with selection on specific genes.

### The effect of the Andean founder event on deleterious alleles

The extreme founder event observed in the Andes could potentially alter genome-wide patterns of deleterious variants beyond the effects of domestication alone. Under a recessive model, maize from the Andes contains significantly more deleterious alleles than any other population (Figure 3; Figure S7; all *p* values < 0.02, Wilcoxon test), and this difference becomes more extreme when considering more severe (*i.e*., higher GERP scores) mutations (Figure S7). In contrast, we observe no significant differences under an additive model (Figure S6; Figure S7). The Andean founder event therefore appears to have resulted in a higher mutation burden of deleterious alleles than seen in other populations of maize. This result is further supported by a higher proportion of fixed deleterious alleles within the Andes and fewer segregating deleterious alleles (Figure S10; Figure 3), a result comparable to the differences observed in load between maize and *parviglumis*.

### Introgression decreases the prevalence of deleterious alleles

Highly variable rates of *mexicana* introgression were detected across our landrace populations (Figure 2; Figure S4; Figure S5). To explore the potential effects of introgression on the genomic distribution of deleterious alleles, we fit a linear model in which the number of deleterious sites is predicted by introgression (represented by 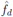) and gene density (exonic base pairs per cM) in 10kb non-overlapping windows in the Mexican highland population where we found the strongest evidence for *mexicana* introgression. Gene density was included as a predictor in the regression to control for the positive correlation observed between gene density and both introgression (*p* = 3.48e — 08) and deleterious alleles (*p* ≈0). When identifying deleterious alleles under both additive and recessive models, we found a strong negative correlation with introgression (*i.e*., fewer deleterious alleles in introgressed regions; *p* ≈ 0 under both models). These findings likely reflect the larger ancestral *N*_*e*_ and more efficient purifying selection in *mexicana*.

## 3 Discussion

Demographic studies in domesticated species have focused largely on identifying progenitor population(s) and quantifying the effect of the domestication bottleneck on genetic diversity [24, 43, 44]. It is likely, however, that the demographic history of domesticates is generally more complex than a simple bottleneck followed by recovery [45, 46]. Many crops and domesticated animals have expanded from defined centers of origin to global distributions, experiencing population size changes and gene flow from closely related taxa throughout their histories [47]. With this in mind, we have characterized maize demography from domestication through initial expansion in order to provide a more complete assessment of the influence of demography on deleterious variants.

### Historical changes in maize population size

Early models of maize demography suggested the ratio of the domestication bottleneck size and duration was between ≈ 2.5: 1 and ≈ 5:1, but little statistical support was found for specific estimates of these individual parameters [23, 28]. Most recently, Beissinger *et al*. [2] fit a model assuming a bottleneck followed by instantaneous exponential recovery. While our results concur with the most recent model in finding a similar bottleneck size (≈ 10% compared to ≈ 5% in Beissinger *et al*.) and that the modern *N*_*e*_ of maize is quite large, the flexibility of MSMC also allowed us to estimate the duration of the bottleneck. We find that the domestication bottleneck may have lasted much longer than previously believed, spanning ≈ 9, 000 generations and only beginning to recover in the recent past (Figure 1B). Analysis of bottlenecks in domesticated African rice and grapes have also suggested a duration of several thousand generations [45, 46], indicating that demographic bottlenecks during crop evolution may have generally occurred over substantial periods of time.

In addition to a strong bottleneck during domestication, our finding that levels of diversity decline in populations increasingly distant from the center of maize domestication are suggestive of serial founder effects during the spread of maize across the Americas (Figure 1C; Figure S1). Serial founder effects are the result of multiple sampling events during which small founder populations are repeatedly drawn from ancestral pools, leading to a stepwise increase in genetic drift and a concomitant decrease in genetic diversity. During maize range expansion, serial founder effects would have occurred if seed carried to each successive colonized region was limited to a sample of whole ears that contained a fraction of the diversity found in the source population [29]. Consistent with serial founder effects, other researchers have found a correlation between geographic and genetic distance in maize landraces [22, 48], though this was previously attributed to limited seed dispersal across the species range leading to isolation by distance (IBD). Neutral expectations of allele frequencies across populations under serial founder effects differ substantially from those predicted under equilibrium conditions [15]. Theory has been developed that predicts the probability that an allele from an origin survives a series of founder effects and reaches high frequency once an expansion is complete [15]. Many of the world’ s crops have experienced such histories of expansion and may be most correctly modeled within this theoretical framework. Studies attempting to identify loci underlying crop adaptation during post-domestication expansion may therefore more accurately compare empirical data to neutral expectations under a serial founder effects demography[15]in order to identify extreme changes in allele frequency driven by selection.

While a history of serial founder effects partially explains the variation in diversity across maize landraces, there are deviations from this model. For example, our combined results showing increased runs of homozygosity (Figure S3), lower nucleotide diversity (Figure S2), and smaller effective population size (Figure 1) in the Andes all suggest a pronounced, ancient founder event and are in agreement with previous work modeling demography in this region [49]. The founder event in the Andes may reflect initially limited cultivation due to the poor performance of maize in this region relative to established root and tuber staples [50]; maize cultivation may have only become widespread after an initial lag period necessary for adaptation. Additionally, we observe somewhat higher than expected nucleotide diversity in maize landraces from the highlands of Mexico and Guatemala (Figure 1C), which may be linked to the introgression we have detected from *mexicana*.

In striking contrast to the bottleneck we observe in maize, the effective population size in *parviglumis* increases steadily from the time of initial maize domestication until the recent past. Multiple population genetic studies have reported negative genome-wide values of Tajima’s D in *parviglumis* from the Balsas River Valley [2, 23, 51], findings characteristic of an expanding population. Likewise, analyses of pollen content in sediment cores from Mexico suggest herbaceous vegetation and grasslands have expanded over the last 10,000 years due to changing environmental conditions during the Holocene and human management of vegetation with fire [52, 53]. While our *parviglumis* samples are drawn from a single population in the Balsas, these data collectively suggest *parviglumis* from this region has experienced expansion over the last several millennia.

Consistent with archaeological evidence of the timing of initial maize domestication[21], we find that maize demographies begin to diverge ≈ 10,000 generations before present, a time that appears to coincide with a steeper decline in maize *N*_*e*_ as well. In contrast, we estimate the timing of the split between maize and our single population of *parviglumis* to be ≈ 75, 000 generations before present, likely reflecting strong population structure in *parviglumis* and divergence of our sampled *parviglumis* from the maize progenitor population. Beissinger *et al*. [2], using samples from additional populations, also find an estimate of maize-*parviglumis* divergence older than the probable onset of domestication, suggesting that currently available sequences of *parviglumis* may not sample well from the populations directly ancestral to domesticated maize.

### The prevalence of gene flow during maize diffusion

Increasingly, range-wide analyses of crops and their wild relatives have identified evidence of gene flow during post-domestication expansion from newly sympatric populations of their progenitor taxa and closely related species [54, 55, 56]. Consistent with previous results from genotyping data [5, 22, 57], we find strong support for introgression from *mexicana* to maize in the highlands of Mexico. While *mexicana* is not currently found in the highlands of Guatemala, we also find strong evidence for *mexicana* introgression in maize from this region, suggesting either *mexicana* was at one time more broadly distributed, or, perhaps more likely, that highland maize from Mexico was introduced to the Guatemalan highlands. Support is also found for *mexicana* introgression in the southwestern US at specific chromosomal regions such as a putative inversion polymorphism on chromosome 3 (Figure 2). These results confirm previous findings suggesting maize from the highlands of Mexico originally colonized the southwestern US [35]. The more limited signal of *mexicana* introgression here may be due to subsequent gene flow from lowland maize as suggested by [35]. Very little evidence is found for *mexicana* haplotypes extending into South America, as highland-adapted haplotypes would likely have been maladaptive and removed by selection as maize traversed the lowland regions of Central America ([49]).

### Impacts of demography on accumulation of deleterious variants

The availability of high-density SNP data from range-wide samples of a species allows for an in-depth assessment of the influence of demography on the prevalence of deleterious alleles. For example, recent studies in both humans and dogs have revealed that historical changes in population size [10, 12, 18] and introgression [9] may contribute substantially to variation in patterns of deleterious variants among populations and across the genome. Previous work in maize has characterized genome-wide trends in deleterious alleles across modern inbred maize lines, revealing that inbreeding during the formation of modern lines has likely purged many recessive deleterious variants [58] and that complementation of deleterious alleles likely underlies the heterosis observed in hybrid breeding programs [58, 59]. Additionally, [2] revealed that purifying selection has removed a greater extent of pairwise diversity (*θ* _π_) near genes in *parviglumis* than in maize due to the higher historical *N*_*e*_ in *parviglumis*, but that this trend is reversed when considering younger alleles due to the recent dramatic expansion in maize population size. To date, however, few links have been made between the historical demography of maize domestication and expansion and the prevalence of deleterious alleles (but see [60] for a comparison of the frequencies of some coding changes). Our analysis reveals that demography has played a pivotal role in determining both the geographic and genomic landscapes of deleterious alleles in maize.

### Population size and deleterious variants

Previous studies have suggested a “cost of domestication” in which a higher burden of deleterious alleles is found in domesticates compared to their wild progenitors [12, 40, 42, 61, 62]. Consistent with these previous studies, we detect an excess of deleterious alleles in maize relative to *parviglumis* (Figure 3; Figure S6; Figure S7), which could be caused by two potential factors. First, reduced population size during the domestication bottleneck could result in deleterious alleles drifting to higher allele frequency. Second, hitchhiking caused by strong positive selection on domestication genes could cause linked deleterious alleles to rise in frequency [12, 61]. While we find support for the former in maize, we see little evidence of the latter. In addition to the cost of domestication, we find a cost of geographic expansion that is likely driven by serial founder effects. The increase in deleterious alleles during expansion is most pronounced in the Andes and may be symptomatic of the extreme founder event we propose above.

Differences in the number of deleterious alleles between maize and *parviglumis* and non-Andean and Andean maize are more dramatic under a recessive model than an additive model. This trend may indicate that the bulk of deleterious alleles in maize are at least partially recessive, such that heterozygous sites contribute less to a reduction in individual fitness. Previous work in human populations has shown that the majority of deleterious mutations are recessive or partially recessive [63], and data from knock-out mutations in yeast have revealed that large-effect mutations tend to be more recessive [64]. Likewise, both theory and empirical evaluation across a number of organisms suggest that mildly deleterious mutations are likely to be partially recessive [65]. In maize, Yang *et al*. [58] have found that most deleterious alleles are at least partially recessive and note a negative correlation between the severity of a deleterious variant and its dominance. Our results thus match nicely both with previous empirical data and theoretical expectations showing that recent population bottlenecks should only show strong differences in load under a recessive model [7].

### Introgression and deleterious variants

Very few studies have investigated the effects of introgression from a taxon with substantially different *N*_*e*_ on the genomic landscape of deleterious variants. The best example is found in the human literature, where confirmation has been found that introgression from Neanderthals with low ancestral *N*_*e*_ increased the overall mutation load in modern humans [9, 19]. We report here the opposite pattern in maize, as introgression appears to have reduced the proportion of deleterious variants. Nonetheless, the underlying interpretation is similar: the introgressing taxon *mexicana* has had a larger ancestral *N*_*e*_ than maize [27], and introgressed haplotypes have thus experienced more efficient long-term purging of deleterious alleles.

## 4 Conclusions

We have demonstrated that demography during the domestication and expansion of maize across the Americas has profoundly influenced putative functional variation across populations and within individual genomes. More generally, we have learned that population size changes and gene flow from close relatives with contrasting effective population size will influence the distribution of deleterious alleles in species undergoing rapid shifts in demography. The significance of our results extends far beyond maize. For example, invasive species that have recently experienced founder events followed by expansion, endangered species subject to precipitous declines in *N*_*e*_, species with a history of postglacial expansion, and new species expanding their range will all likely show clear genetic signals of the interplay between demography and selection. This interaction bears importantly on the adaptive potential of both individual populations and species. By fully characterizing this relationship we can better understand how the current evolutionary trajectory of a species has been influenced by its history.

## 5 Materials and Methods

### Samples, whole genome resequencing, and read mapping

A total of 31 maize landrace accessions were obtained from the US Department of Agriculture (USDA)’s National Plant Germplasm System and through collaborators (Figure S12). Samples were chosen from four highland populations (Andes, Mexican Highlands, Guatemalan Highlands and southwestern US Highlands) and two lowland populations (Mexican and South American lowlands) (Figure 1A). In addition, four open-pollinated *parviglumis* samples were selected from a single population in the Balsas River Valley in Mexico. DNA was extracted from leaves using a standard cetyltrimethyl ammonium bromide (CTAB) protocol [66]. Library preparation and Illumina HiSeq 2000 sequencing (100-bp paired-end) were conducted by BGI following their established protocols. BWA v. 0.7.5.a [67] was used to map reads to the maize B73 reference genome v3 [68] with default settings. The duplicate molecules in the realigned bam files were removed with MarkDuplicates in Picardtools v. 1.106 (http://broadinstitute.github.io/picard) and indels were realigned with GATK v. 3.3-0 [69]. Sites with mapping quality less than 30 and base quality less than 20 were removed and only uniquely mapped reads were included in downstream analyses.

### Demography of maize domestication and diffusion

The recently developed method MSMC [31] was utilized to infer effective population size changes in both *parviglumis* and maize. SNPs were called via HaplotypeCaller and filtered via VariantFiltration in GATK [69] across all samples. SNPs with the following metrics were excluded from the analysis: QD < 2.0; FS > 60.0; MQ < 40.0; MQRankSum <-12.5ReadPosRankSum <-8.0. Vcftools v. 0.1.12 [70] was used to further filter SNPs to include only bi-allelic sites. SNPs were phased using BEAGLE v. 4.0 [71] with SNP data from the maize HapMap2 panel [60] used as a reference. Only sites with depth between half and twice of the mean depth were included in analyses. In addition, the software SNPable (http://lh3lh3.users.sourceforge.net/snpable.shtml) was used to mask genomic regions in which reads were not uniquely mapped. The mappability mask file for MSMC was generated by stepping in 1 *bp* increments across the maize genome to generate 100 *bp* single-end reads, which were then mapped back to the maize B73 reference genome [68]. Sites with the majority of overlapping 100-mers mapped uniquely without mismatch were determined to be “SNPable” sites and used for the MSMC analyses. For effective population size inference in MSMC, we used 5 × 4 + 25 × 2 + 5 × 4 as the pattern parameter and the value *m* was set as half of the heterozygosity in *parviglumis* and maize populations, respectively.

In order to explore the trend of genetic diversity away from the domestication center, the correlation between the percentage of polymorphic sites and the geographic distance to the Balsas River Valley (latitude: 18.099138; longitude: -100.243303) was examined via linear regression. Geographical distance in kilometers was calculated based on great circle distance using the haversine transformation [17].

### Population structure, genetic diversity and inbreeding coefficients

We first evaluated population structure by principal components analysis (PCA) with ngsCovar [72] in ngsTools [73] based on the matrix of posterior probabilities of SNP genotypes produced in ANGSD v. 0.614 [74], and then utilized NGSadmix v. 32 [75] to investigate the admixture proportion of each accession. The NGSadmix analysis was conducted based on genotype likelihoods for all individuals, which were generated with ANGSD (options-GL2-doGlf 2-SNP_pval 1*e* — 6), and K from 2 to 10 was set to run the analysis for sites present in a minimum of 77 % of all individuals (24 in 31). A clear outlier in the Mexican Highland population was detected, RIMMA0677, a sample from relatively low altitude, which was suspected to contain a divergent haplotype. A neighbor joining tree of SNPs within an inversion polymorphism on chromosome 4 that includes a diagnostic highland haplotype [5] was constructed with the R package phangorn [76]. The sample RIMMA0677 was not clustered with other highland samples, but embedded within lowland haplotypes (Figure S11), so it was removed from further analyses.

The genetic diversity measures Watterson’s *θ* and *θ*π were calculated in ANGSD [74] for each population. The neutrality test statistic Tajima’s D was calculated with an empirical Bayes approach [77] implemented in ANGSD by first estimating a global site frequency spectrum (SFS) then calculating posterior sample allele frequencies using the global SFS as a prior. The three statistics were summarized across the genome using 10-kb non-overlapping sliding windows.

Inbreeding coefficients for each individual were estimated with ngsF [78] with initial values of set to be uniform at 0.01 with an epsilon value of 1*e*—5.

In addition, SNPs were polarized using the *T*. *dactyloides* genome to *assess* the frequency of derived homozygous sites in each maize landrace population. T. *dactyloides* short reads were downloaded from the NCBI SRA database (SRR447804 - SRR447807), mapped to the B73 reference genome v3 ([68]) with BWA [67] and incorporated into SNP calling as described above.

### Runs of Homozygosity

SNPs were down-sampled to contain one SNP in a 2-*kb* window to identify segments representing homozygosity by descent (*i.e*., autozygosity) rather than by chance. PLINK v. 1.07 (http://pngu.mgh.harvard.edu/purcell/plink/) [79] was applied to identify segments of ROH in a window containing 20 SNPs, among which the number of the maximum missing SNPs was set to 2 and the number of the maximum heterozygous sites was set to 1. The shortest length of final ROHs was set to be 300 *kb*. Physical distances were converted into genetic distances based on a recent genetic map [80].

### Detection of Introgression

To assess per-genome evidence of population admixture between maize landraces and teosinte, we calculated the D statistic using ANGSD [74]. The statistic was calculated using trees of the form (((X, low),*mexicana*),T. *dactyloides*). One accession from the Mexican Lowland population was randomly sampled as the “low” taxon, and each sample from all other populations except the Mexican Lowland was set as “X”. The mexicana accession TIL25 from the maize HapMap2 project [60] was treated as the third column species. The D statistic was calculated in a 1*kb*-block and then jackknife bootstrapping was conducted to estimate significance.

In addition, the 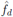 statistic [36] was calculated based on a similar tree form (((*P*_1_, *P*_2_), *P*_3_), O), but using allele frequencies across multiple individuals for each position on the tree. We fixed *P*_1_ as the Mexican Lowland population, *P*_3_ as two lines of *mexicana* (TIL08 andTIL25) and T. *dactyloides* as the outgroup. *P*_2_ was set to each of the four highland populations and the South American Lowland population.

The 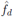 statistic was calculated in 10-*kb* non-overlapping windows across the genome with the python script egglib_sliding_windows.py (https://github.com/johnomics/Martin_Davey_Jiggins_evaluating_introgression_statistics), which makes use of the EggLib library [81]. The input file was generated by first identifying genotypes using ANGSD (- doMajorMinor 1-doMaf 1-GL 2-doGeno 4-doPost 1-postCutoff 0.95 -SNP_pval 1*e* — 6) followed by format adjustments with a custom script (Please see “Data and analysis pipeline accessibility”).

### Estimating burden of deleterious mutations

We estimated the individual burden of deleterious alleles based on GERP scores [82] for each site in the maize genome, which reflects the strength of purifying selection based on constraint in a whole genome alignment of 13 plant species [83]. The alignment and estimated GERP scores are available at iplant (/iplant/home/lilepisorus/GERPv3).Scores above 0 may be interpreted as historically subject to purifying selection, and mutations at such sites are likely deleterious. We identified *Sorghum* bicolor alleles in the multiple species alignment as ancestral and defined the non-*Sorghum* allele as the deleterious allele. Only biallelic sites were included for our evaluation. Inclusion of the maize B73 reference genome when calculating GERP scores [83] introduces a bias toward underestimation of the burden of deleterious alleles in maize versus teosinte populations.Therefore, we corrected the GERP scores of sites where the B73 allele is derived following [7]. Briefly, we divided SNPs where the B73 allele is ancestral into bins of 1% derived allele frequency based on maize HapMap3 [84] and used this frequency distribution to estimate the posterior probability of GERP scores for SNPs where the B73 allele is derived.

The sum of GERP scores multiplied by deleterious allele frequency for each SNP site was used as a proxy of individual burden of deleterious alleles under an additive model (*HET* * 0.5 + *HOM* * 1). This burden was calculated under a recessive model as the sum of GERP scores multiplied by one for each deleterious homozygous site (*HOM* * 1). For a better understanding of the variation of individual burden among sites under varied selection strength, we partitioned the deleterious SNPs into four categories (-2 <GERP≤ 0, nearly neutral; 0 < GERP ≤ 2, slightly deleterious; 2 < GERP ≤ 4, moderately deleterious; GERP *>* 4, strongly deleterious) and recapitulated the above statistics.

### Data and analysis pipeline accessibility

The pipeline and custom scripts utilized in this paper are documented in the following GitHub repository: https://github.com/HuffordLab/Wang_Private/tree/master/demography/ analyses The WGS raw reads have been deposited in NCBI SRA (SRP065483).

## 6 Acknowledgments

This study was supported by the US Department of Agriculture (USDA #2009-65300-05668), the USDA Agricultural Research Service, the National Science Foundation (NSF IOS #1546719), USDA Hatch project (CA-D-PLS-2066-H), and startup funds from Iowa State University. Additionally, we thank Dr. Andrew Severin and Dr. Arun Seetharam for bioinformatic support.

## Supporting Information

**Figure S1:**
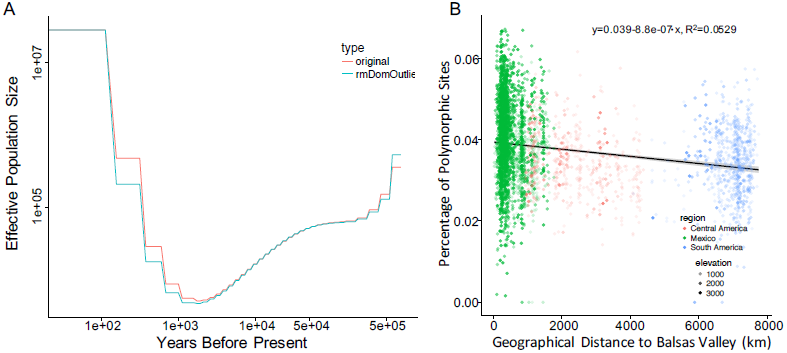
Demography of maize populations. A. MSMC results before and after masking candidate regions under selection during domestication. B. Percentage of heterozygous sites versus distance from the Balsas Valley in 3520 samples from the SeeDs data set.

**Figure S2:**
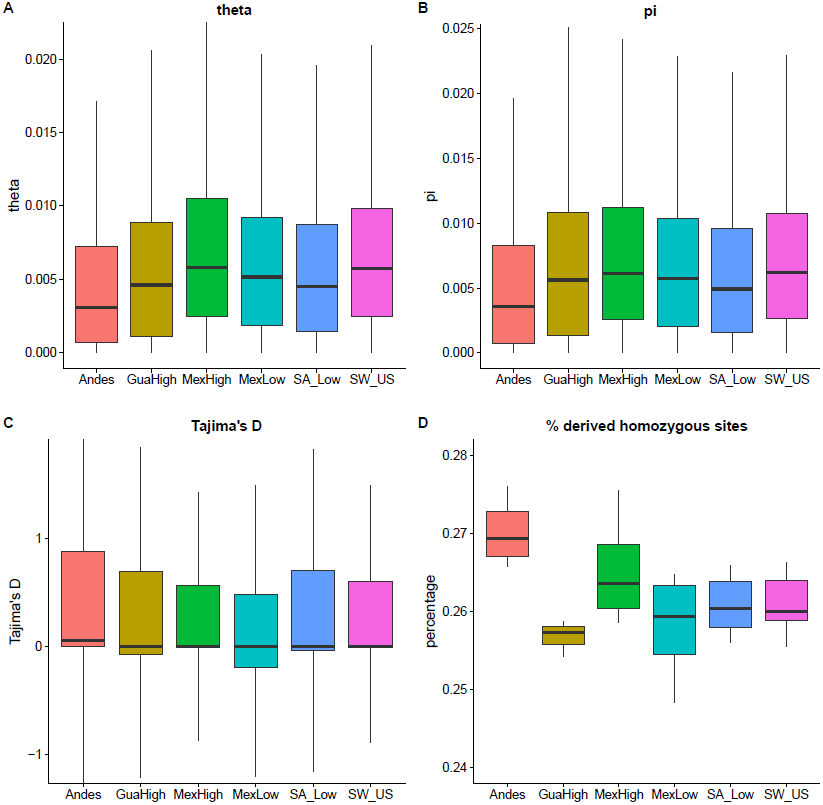
Boxplot of multiple population genetic statistics. Watterson’s *theta* (A), *θ*_π_(B) and Tajima’s D (C) are based on values in 10-kb non-overlapping windows across the genome. Percentage of derived homozygous sites was calculated for each individual and reported per population

**Figure S3:**
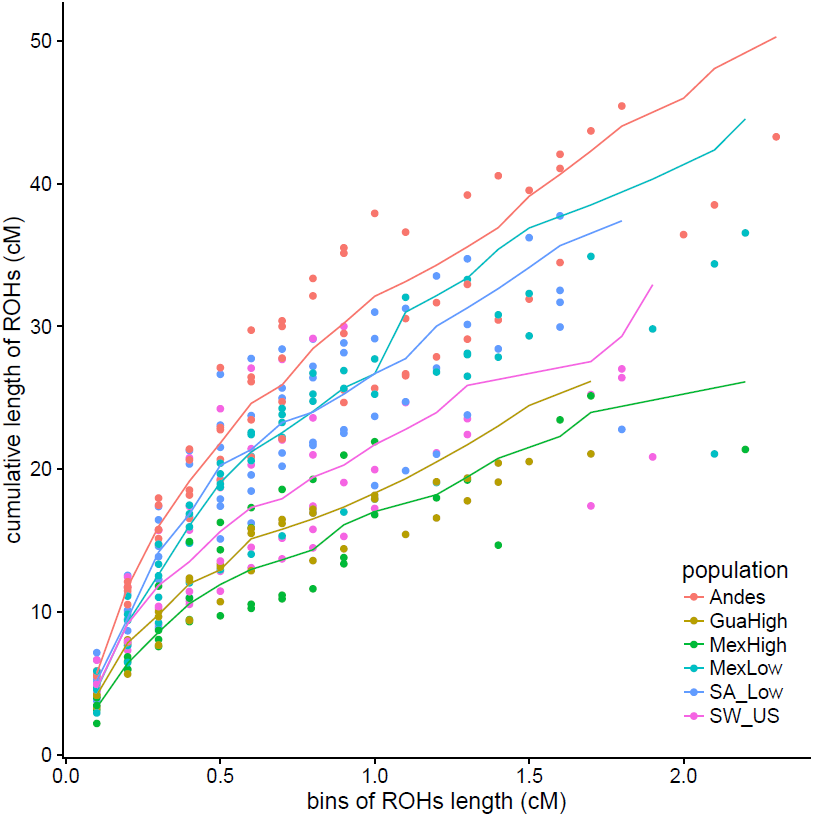
Cumulative length of ROHs in cM among populations. The lines indicate median level in each population. ROH: runs of homozygosity.

**Figure S4:**
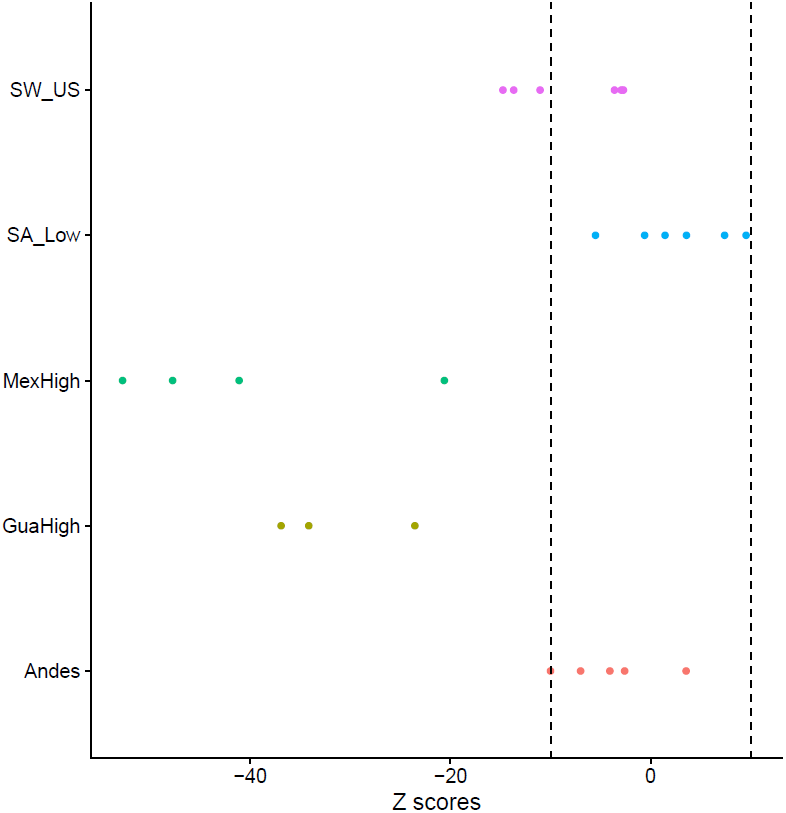
Introgression from *mexicana* into maize landraces. Evidence of introgression from *mexicana* into Mexican highland, Guatemalan highland and Southwestern US highland maize populations. The dashed lines correspond to *Z* scores equal to -10 and 10.

**Figure S5:**
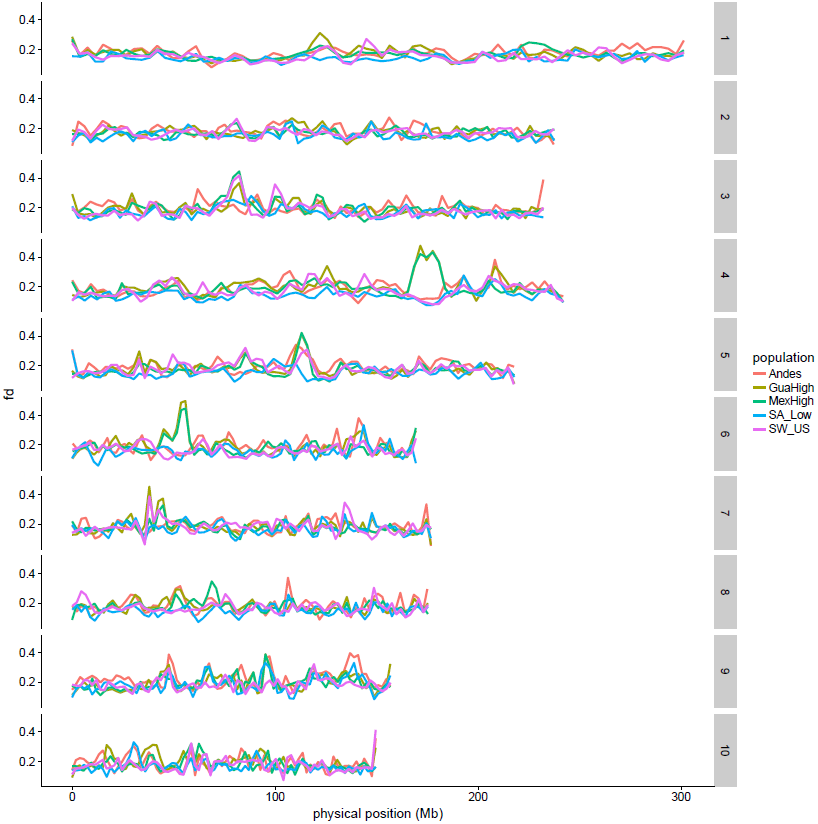
Loess regression of 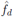 in 10-kb nonoverlapping windows across all chromosomes.

**Figure S6:**
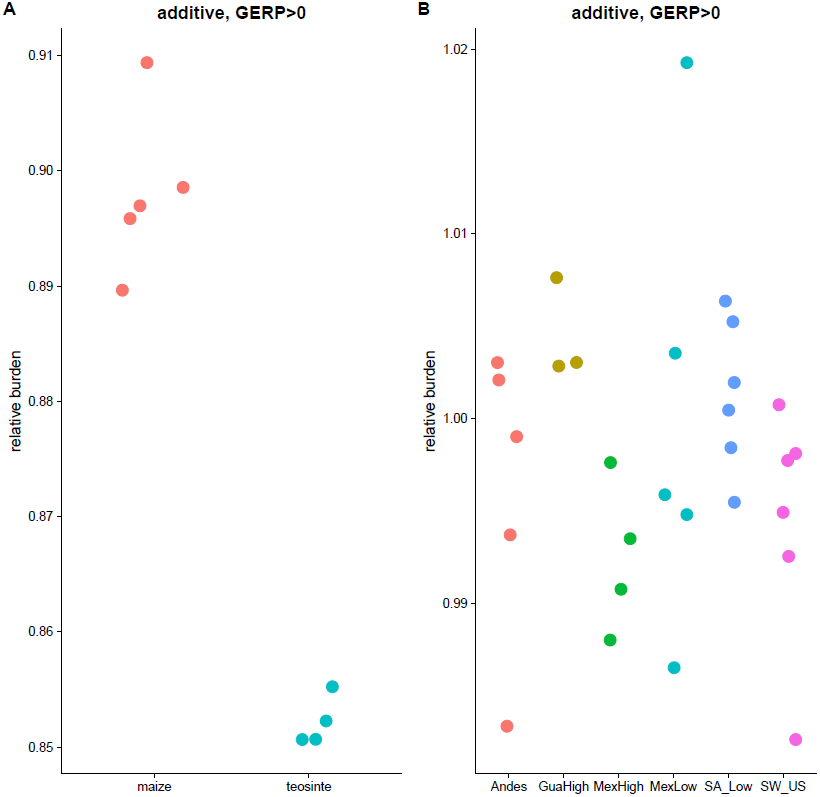
Relative burden of deleterious alleles under additive model between maize and teosinte (A) and among maize populations (B).

**Figure S7:**
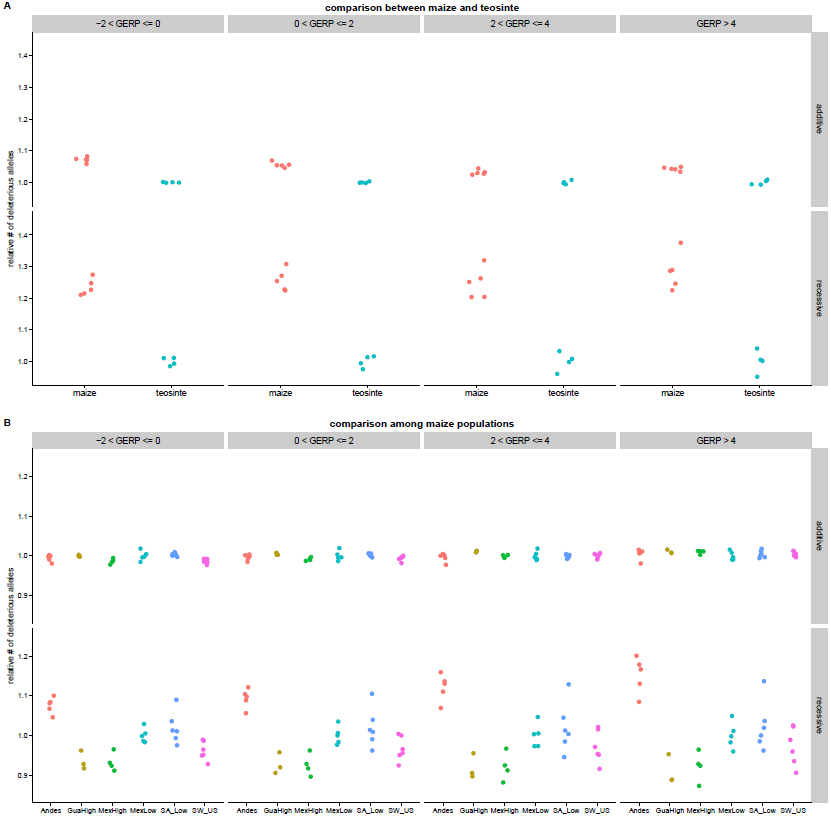
Relative burden of deleterious alleles under both additive and recessive models with different GERP partitions between maize and teosinte (A) and among maize populations (B).

**Figure S8:**
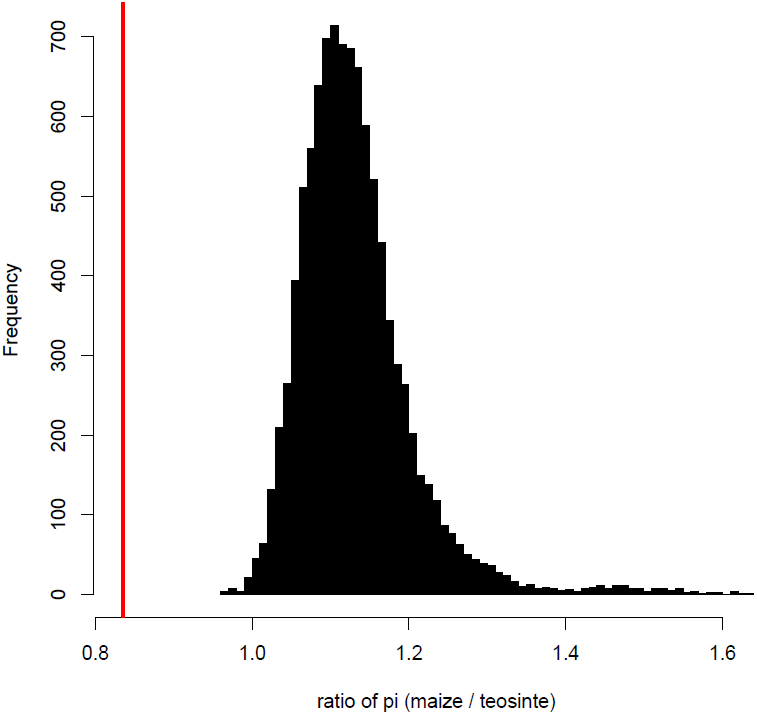
Distribution of ratio of *θ*_π_ between maize and teosinte in 420 domestication candidate genes (mean value was indicated with red line) compared to 10,000 replicates of genome-wide sampling of 420 random genes.

**Figure S9:**
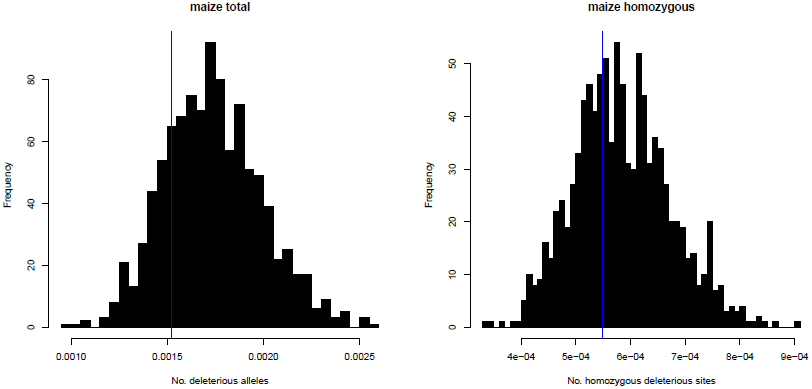
Distribution of number of deleterious sites per bp in 420 domestication candidate genes (indicated with blue line) compared to genome-wide random samples under an (A) additive model and (B) recessive model.

**Figure S10:**
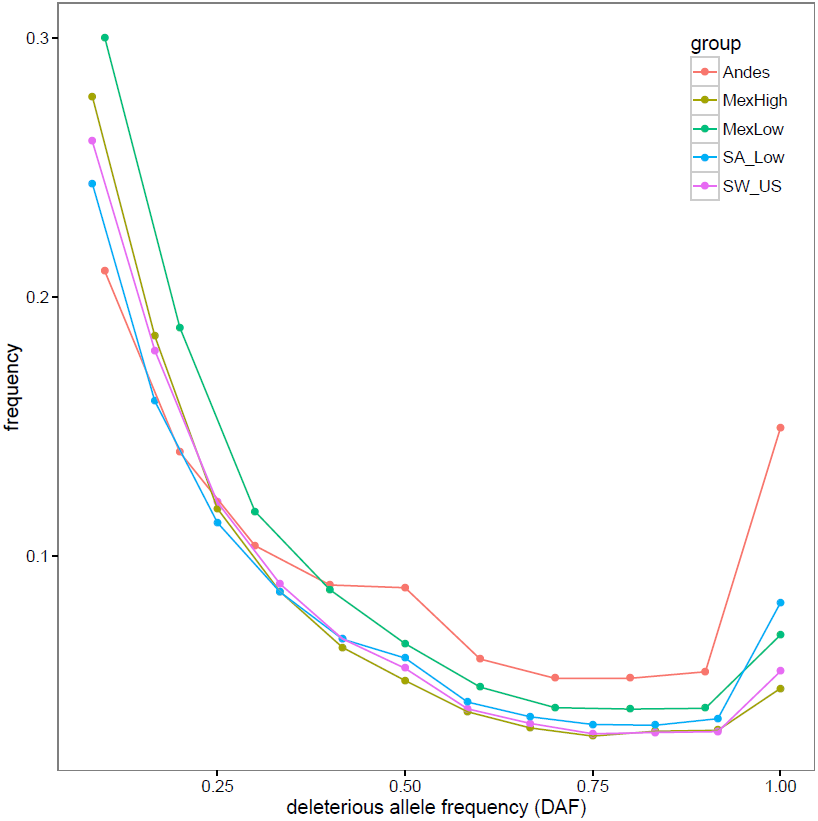
Site frequency spectrum of deleterious SNPs in five populations; GuaHigh is not included since the small sampling limited power for the SFS.

**Figure S11:**
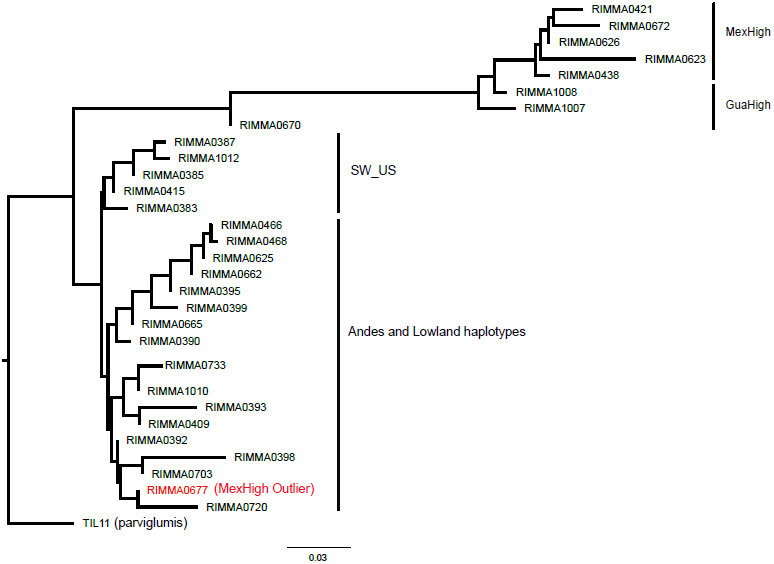
Neighbor Joining tree of SNPs from an inversion on chromosome 4 with a diagnostic haplotype for highland Mexican material.

**Figure S12:**
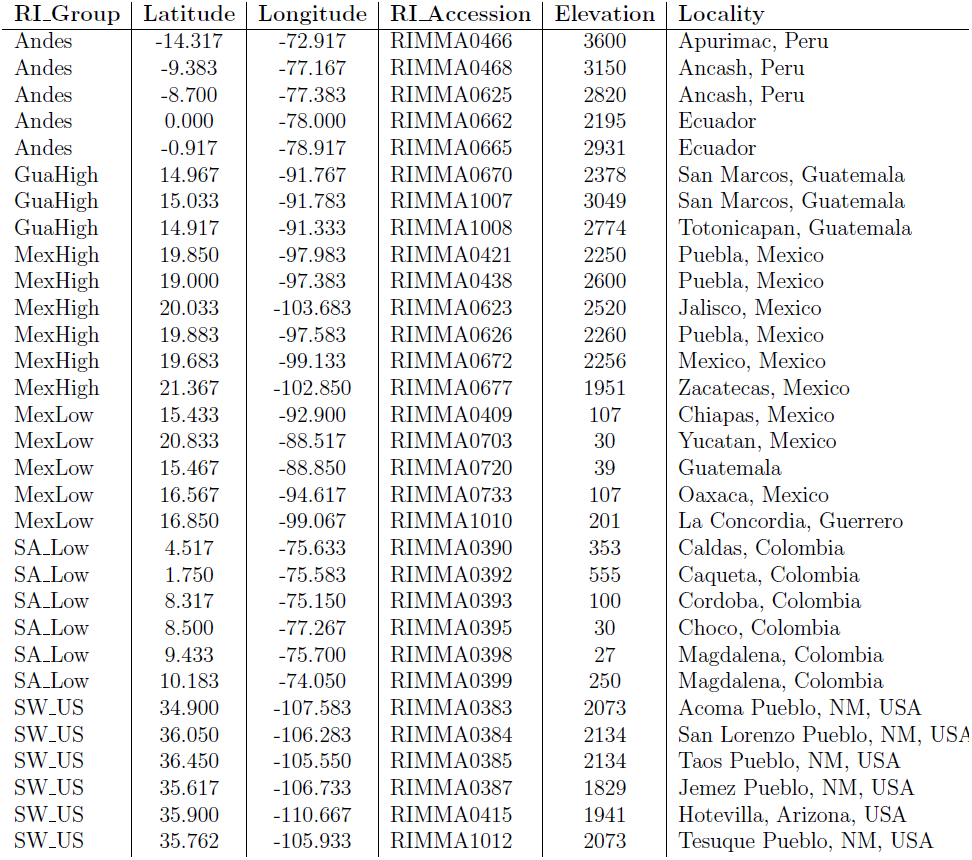
Basic information regarding the sampled maize landrace accessions. NM: New Mexico.

